# NK cell metabolic adaptation to infection promotes survival and viral clearance

**DOI:** 10.1101/2021.03.17.435833

**Authors:** Francisco Victorino, Tarin Bigley, Eugene Park, Cong-Hui Yao, Jeanne Benoit, Li-Ping Yang, Sytse J Piersma, Elvin J Lauron, Rebecca M Davidson, Gary J Patti, Wayne M Yokoyama

## Abstract

Natural killer (NK) cells are essential for early protection against virus infection, and must metabolically adapt to the energy demands of activation. Here, we found upregulation of the metabolic adaptor hypoxia inducible factor-1α (HIF-1α) is a feature of NK cells during murine cytomegalovirus (MCMV) infection *in vivo*. HIF-1α-deficient NK cells failed to control viral load, causing increased morbidity. No defects were found in effector functions of HIF-1α KO NK cells however, their numbers were significantly reduced. Loss of HIF-1α did not affect NK cell proliferation during *in vivo* infection and *in vitro* cytokine stimulation. Instead, we found HIF1α-deficient NK cells showed increased expression of the pro-apoptotic protein Bim and glucose metabolism was impaired during cytokine stimulation *in vitro*. Similarly, during MCMV infection HIF1α-deficient NK cells upregulated Bim and had increased caspase activity. Thus, NK cells require HIF-1α-dependent metabolic functions to repress Bim expression and sustain cell numbers for an optimal virus response.

## Introduction

During infection with a cytolytic virus, such as murine cytomegalovirus (MCMV), infected cells become sources of viral replication and of target cells. MCMV elicits non-specific NK cell activation via inducing a strong ultimately succumb due to direct viral cytopathology. Natural killer (NK) cells play an integral role in initiating and fine-tuning immune responses to provide protection against MCMV. NK cells carry out this role through two overlapping functional properties: 1) secretion of pro-inflammatory cytokines; and 2) direct killing host cytokine response and also triggers specific NK cell activation in C57BL/6 mice. In particular, the Ly49H activation receptor recognizes the viral encoded protein m157 on infected cells, triggering cytotoxicity and cytokine production that can modulate subsequent immune responses [1-5]. These non-specific and specific NK cell responses are integrated to provide optimal NK cell responses and protection [6]. Ly49H^+^ NK cells subsequently acquire properties characteristic of adaptive immunity of expansion, contraction, and generation of memory-like responses [7]. Thus, NK cells function as sentries during MCMV infection, controlling pathogen propagation and inflammation through their effector functions.

During inflammation, rapid energy consumption by cells for activation, biosynthesis, and the generation of reactive oxygen species results in hypoxia, a reduction in oxygen levels, even in the presence of environmental normoxia. Hypoxia-inducible factor 1-alpha (HIF1α) is an evolutionarily conserved transcription factor whose protein expression is determined in an oxygen-dependent manner [8-11]. Briefly, when oxygen is readily available, hydroxylation of target residues on HIF1α results in ubiquitination and proteasome degradation while, during hypoxia this modification does not occur, allowing for protein accumulation and translocation into the nucleus. Under hypoxic conditions, HIF1α targets genes that contain hypoxia-response elements (HREs) to initiate transcriptional changes for adaption, supporting multiple cellular processes such as cell proliferation, survival, and glucose metabolism [12, 13]. Thus, HIF1α expression is directly associated with degree of oxygen levels and is necessary for adaption to hypoxia during inflammation.

Hypoxia is a fundamental characteristic of tumor environments and drives HIF1α protein expression *in situ*. HIF1α has been considered a potential mediator of suppressive effects in NK cells which demonstrate impaired effector functions in hypoxia *in vitro* that mirror tumor-infiltrating NK cells [14-16]. Indeed, deletion of HIF1α in NK cells resulted in an improved ability to reduce tumor burden in two independent studies which was attributed to a dysfunctional VEGF axis or increased effector functions through IL-18 signaling [17, 18]. HIF1α also supports metabolic adaptation to the tumor environment by upregulating key glycolysis genes to support cell proliferation and survival. Interestingly, Ni et al. showed HIF1α KO NK cells had higher oxidative phosphorylation (OXPHOS) after cytokine activation, presumably to compensate for poor glucose metabolism in the absence of HIF1α upregulated genes. However, measurements of glucose metabolism showed no differences compared to control mice. Using similar *in vitro* cytokine stimulation experiments, another study showed that glucose metabolism in NK cells was regulated by cMyc, and HIF1α was found to be dispensable and the absence of HIF1α had no effect on OXPHOS metabolism [19]. Regardless, as a caveat, the pro-inflammatory cytokines used are more associated with pathogen infection rather than a tumor environment. Therefore, the contribution of HIF1α to NK cell metabolism is incompletely understood, particularly during the pathogen infection.

In this paper, we utilized *in vitro* assays and MCMV infections to study the role of HIF1α in NK cells. Compared to control mice, the absence of HIF1α in NK cells resulted in susceptibility to infection but this was not due to an impaired effector response. Instead, we show a previously unknown role for HIF1α in NK cells during MCMV infection, supporting survival rather than proliferation through efficient glucose metabolism that is required for an optimal host response to virus infection.

## Results

### HIF1α is basally expressed in NK cells and is upregulated by MCMV infection

We found HIF1α protein was readily detectable in splenic NK cells from naïve mice, immediately *ex vivo* using anti-HIF1a (**Figure 1A**). By contrast, there was essentially no HIF1α expression in other naïve lymphocytes directly *ex vivo* (**Figure 1B**). Under hypoxic conditions as compared to normoxia, HIF1α protein expression increased in NK cells cultured *in vitro* (**Figure 1C**). Interestingly, under normoxic conditions, IFNβ + IL18 upregulated HIF1α expression to levels comparable to hypoxic conditions, though these cytokines did not further increase HIF1α expression under hypoxic conditions. Moreover, when we cultured splenocytes in IL-15 under normoxic conditions for 16, 24, and 48 hours, we found highest expression at 48 hours (**Figure 1D**). Thus, NK cells constitutively express HIF1α immediately *ex vivo* that can be increased by hypoxia alone, or cytokine stimulation in an oxygen-rich environment.

**Figure 1.**
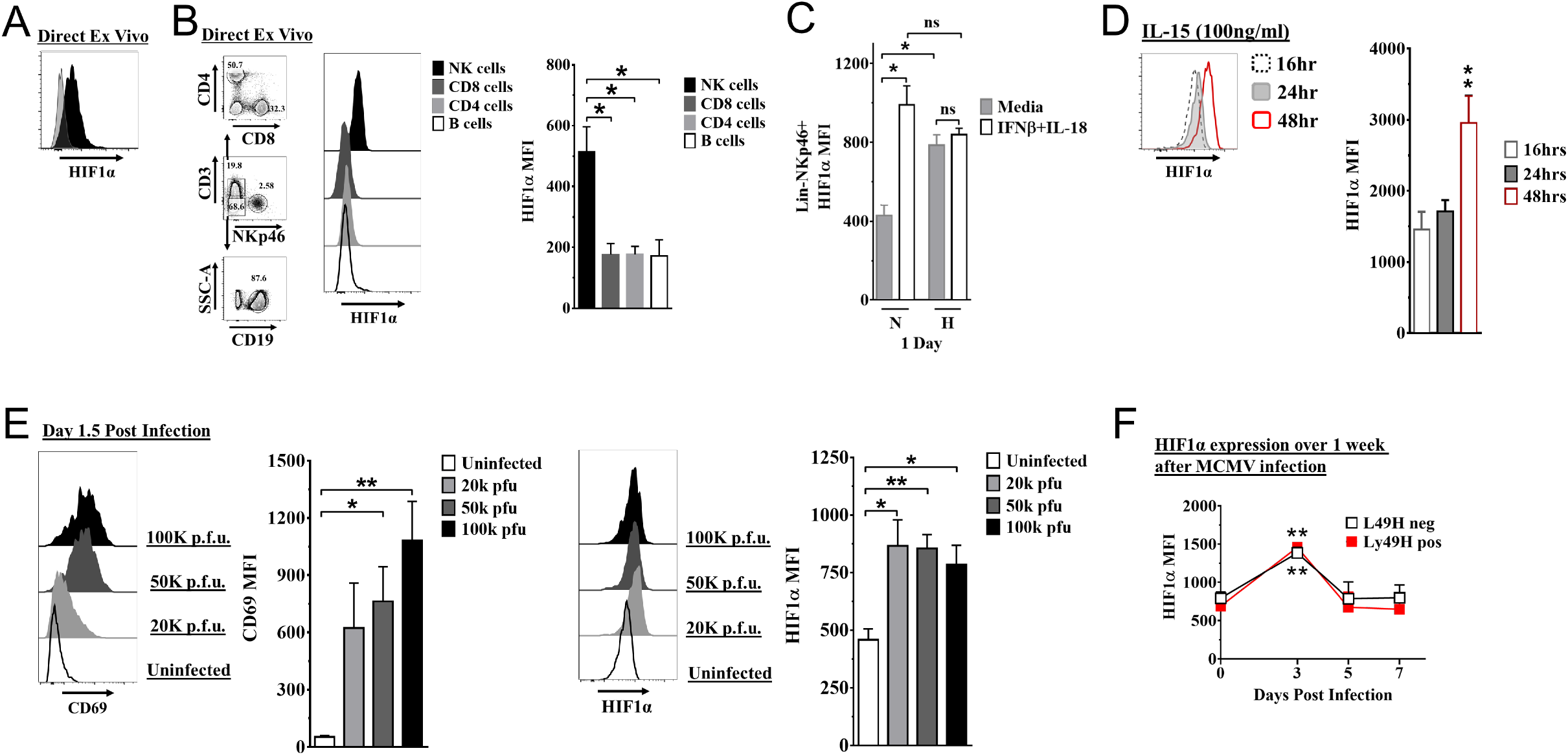
HIF1α expression in NK cells. (A) HIF1α protein expression in splenic NK cells (black) *ex vivo* compared to FMO control (gray) by flow cytometry. Representative experiment of two independent experiments. (B) Comparison of lymphocytes HIF1α protein expression. Bar graphs show data from 2 independent experiments with 4 mice per group. (C) Spleens from C57BL/6 mice were prepared as in methods and were stimulated with IFNβ (200units) +IL-18 (50ηg/ml) for 24 hours in normoxia (N, 20%) or hypoxia (H, 1%). HIF1α expression in NK cells was determined. Data are from 2 independent experiments with 3 mice per group. (D) Splenic NK cells were cultured in 25ηg/ml IL-15 for 48 hours and HIF1α expression was measured at 16, 24, and 48 hours. Data are from 2 independent experiments with 4 mice per group. (E) CD69 and HIF1α was determined at day 1.5 pi. Data are from 2-3 independent experiments with 3 mice per group. Statistical significance for uninfected versus indicated dose. (F) HIF1α expression in Ly49H^+^ or Ly49H^−^ NK cells over 7 days from C57BL/6 mice that were infected with 50k p.f.u. MCMV. Statistical significance for d0 versus indicated day. Data are from 2 independent experiments with 3-7 mice per group. All data depict mean +/− SEM, with each data set containing data indicated number of mice per group from independent experiments. Unpaired t-test was performed (A, B, D-F) or 1-way ANOVA (C). Statistical significance indicated by n.s., no significant difference; * = *p*<.05; ** = *p*<.01; *** = *p*<.001; **** = *p*<.0001.

To determine HIF1α expression in NK cells during an immune response *in vivo*, we studied C57BL/6 mice infected with MCMV. At all doses of MCMV, splenic NK cell activation was observed on day 1.5 post-infection (pi) as indicated by CD69 expression, an activation marker not expressed on naïve NK cells (**Figure 1E**), and concomitant with increased HIF1α expression. HIF1α expression was highest at day 3 pi regardless of expression of the Ly49H receptor which activates NK cell proliferation and memory formation (**Figure 1F**). Thus, MCMV infection drives upregulation of HIF1α expression in NK cells, independent of activation receptor stimulation.

### NK cells require HIF1α for an optimal response to virus infection

To determine if HIF1α may contribute to NK cell function *in vivo*, we crossed NKp46-iCre mice with HIF1α-Flox mice for NK cell-specific deletion of HIF1α (1αKO) for comparison to HIF1α-Flox (FL) control mice which do not express Cre. The frequencies and numbers of NK cells in 1αKO mice were comparable to FL controls (**Figure S1A**) and there was normal NK cell development, as indicated by CD27 and CD11b expression profiles (**Figure S1B**). Expression of other functional markers and receptors, such as DNAM-1, KLRG1, NKG2A (**Figure S1C**), and the Ly49 repertoire, was also normal (**Figure S1D**). These data support previous findings that HIF1α is dispensable for normal NK cell development [17].

During MCMV infection, however, we found 1αKO mice showed increased weight loss as compared to control FL mice (**Figure 2A**). This increased morbidity was coupled with an impaired ability to control viral load at day 5 pi (**Figure 2B**). The number of NK cells in 1α KO mice declined at day 1.5 pi and was significantly reduced at day 3 pi (**Figure 2C**), consistent with the kinetics of HIF1α expression in infected WT mice (**Figure 1C, D**). While FL control mice showed the characteristic expansion and contraction of Ly49H^+^ NK cells over 1 month of infection as previously reported [7], the 1αKO mice displayed reduced Ly49H^+^ NK cell expansion (**Figure 2D**). Thus, during MCMV infection, NK cells require HIF1α to protect against morbidity, control viral load, sustain NK cell numbers, and support Ly49H^+^ NK cell expansion.

**Figure 2.**
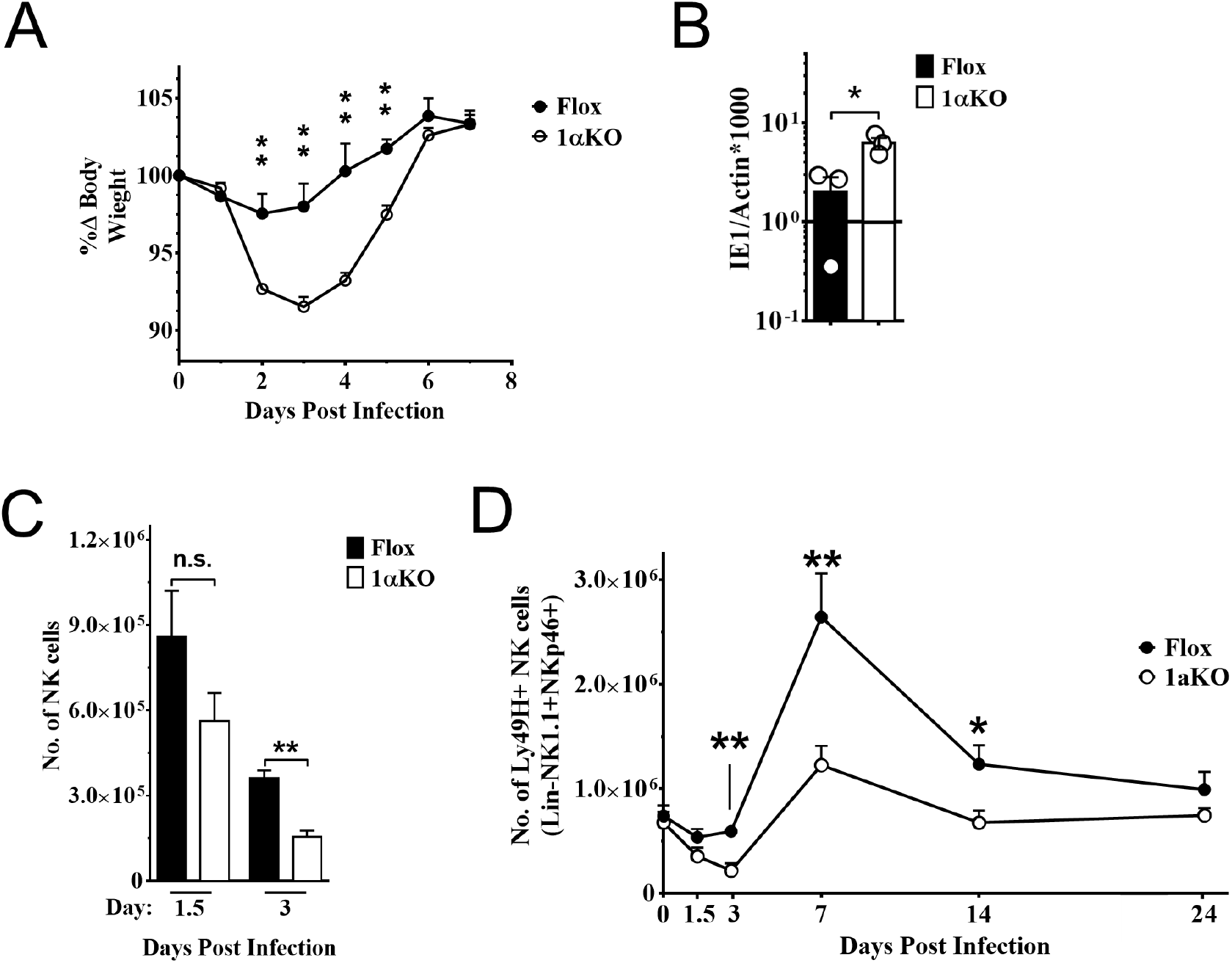
NK cells require HIF1α for an optimal response to virus infection. (A) 1αKO or FL control mice were infected with 50K p.f.u. then monitored for weight loss over seven days. Each data represents four mice from two independent experiments. (B) Splenic viral load in 1αKO or FL control mice were infected with 50K p.f.u. then spleens harvested day 5 post infection. Data are from 3 independent experiments with 3 mice per group. (C) Number of bulk NK cells was quantified at days 1.5 and 3 pi from 1αKO or FL control mice. Data are from 2-3 independent experiments with 4-6 mice per group. (D) 1αKO or FL control mice were infected with 50K p.f.u. and Ly49H^+^ NK cell expansion was determined at day 1.5, 3, 7, 14, and 24 days pi. Data are from 3 independent experiments with 4-9 mice per group. Data depict mean +/− SEM, with each data set containing data indicated number of mice per group from independent experiments. Unpaired t-test was performed on A-D. Statistical significance indicated by n.s., no significant difference; * = *p*<.05; ** = *p*<.01; *** = *p*<.001; **** = *p*<.0001.

### HIF1α is dispensable for NK cell effector function

To determine how HIF1α is required by NK cells to provide immunity during MCMV infection, we assessed if HIF1α contributed to NK cell activation and effector function. Surprisingly, we found that 1αKO NK cells at day 1.5 pi had significant increases in CD69 expression, and IFNγ production with decreased KLRG1 expression as compared to uninfected 1αKO NK cells (**Figure 3A**). While these changes were subtly more than infected FL NK cells, such changes were not readily apparent at day 3 pi (**Figure 3B**). Thus, HIF1α is not required for general activation of NK cells during MCMV infection and may actually restrict NK activation early in infection.

**Figure 3.**
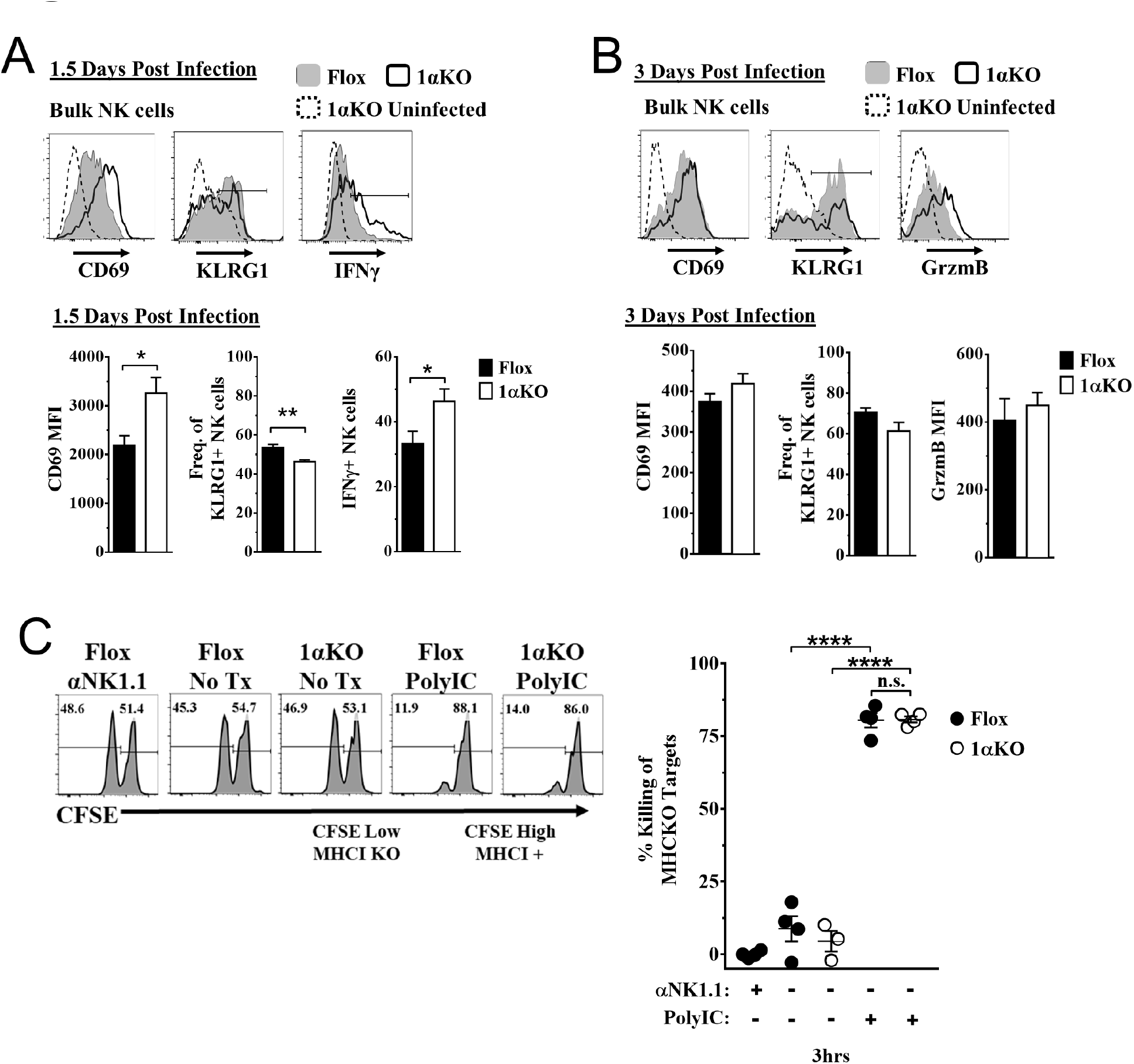
HIF1α is dispensable for NK cell effector function. (A) Representative histograms (top) and quantification (bottom) of CD69 MFI on NK cells and frequency of KLRG1^+^ and IFNγ^+^ NK cells at day 1.5 pi of 1αKO or FL control mice. Data are from 2-3 independent experiments with 4-6 mice per group. (B) Representative histograms (top) and quantification (bottom) of CD69 and Granzyme B MFI (GrzmB) of NK cells, and frequency of KLRG1^+^ NK cells at day 3 pi. Data are from 2 independent experiments with 4 mice per group. (C) 1αKO or FL control mice were injected with PBS or Poly (I:C) then 3 days later 20×10^6^ splenocytes at 1:1 ratio of CFSE-labeled MHCI-sufficient (CFSE^High^) and -deficient (CFSE^Low^) were intravenously transferred into mice. MHC-deficient splenocyte elimination was measured 3 hours post-transfer by flow cytometry. Data are from 3 independent experiments with 3-4 mice per group. Data depict mean +/− SEM, with each data set containing data indicated number of mice per group from independent experiments. Unpaired t-test was performed on (A &B) or 1-way ANOVA (C). Statistical significance indicated by n.s., no significant difference; * = *p*<.05; ** = *p*<.01; *** = *p*<.001; **** = *p*<.0001.

To directly assess HIF1α-dependent NK cytolytic capacity *in vivo* without the confounding issue of decreased NK cell numbers during MCMV infection in 1αKO mice, we performed missing-self rejection assays in uninfected mice after TLR stimulation with poly-I:C. Both cohorts of mice were equally efficient killers of MHCI-deficient targets (**Figure 3C**). Thus, *in vivo* cytotoxic killing function is intact in HIF1α KO NK cells, suggesting that the impaired NK cell immunity to MCMV is not related to inability to kill target cells.

### HIF1α KO NK cells divide normally but numbers are reduced

Another possible explanation for the impaired NK cell immunity to MCMV observed in 1α KO mice could be a defect in NK cell proliferation. However, we observed a modest increase in the expression of proliferation marker Ki67 in bulk NK cells at day 1.5 pi and higher expression at day 3 pi that were comparable between infected 1α KO and control FL mice (**Figure 4A**). Similar results were seen with BrdU incorporation at day 3 (**Figure S2A**), confirming that HIF1α is not required during the non-specific phase of NK cell proliferation that occurs early after MCMV infection [6].

**Figure 4.**
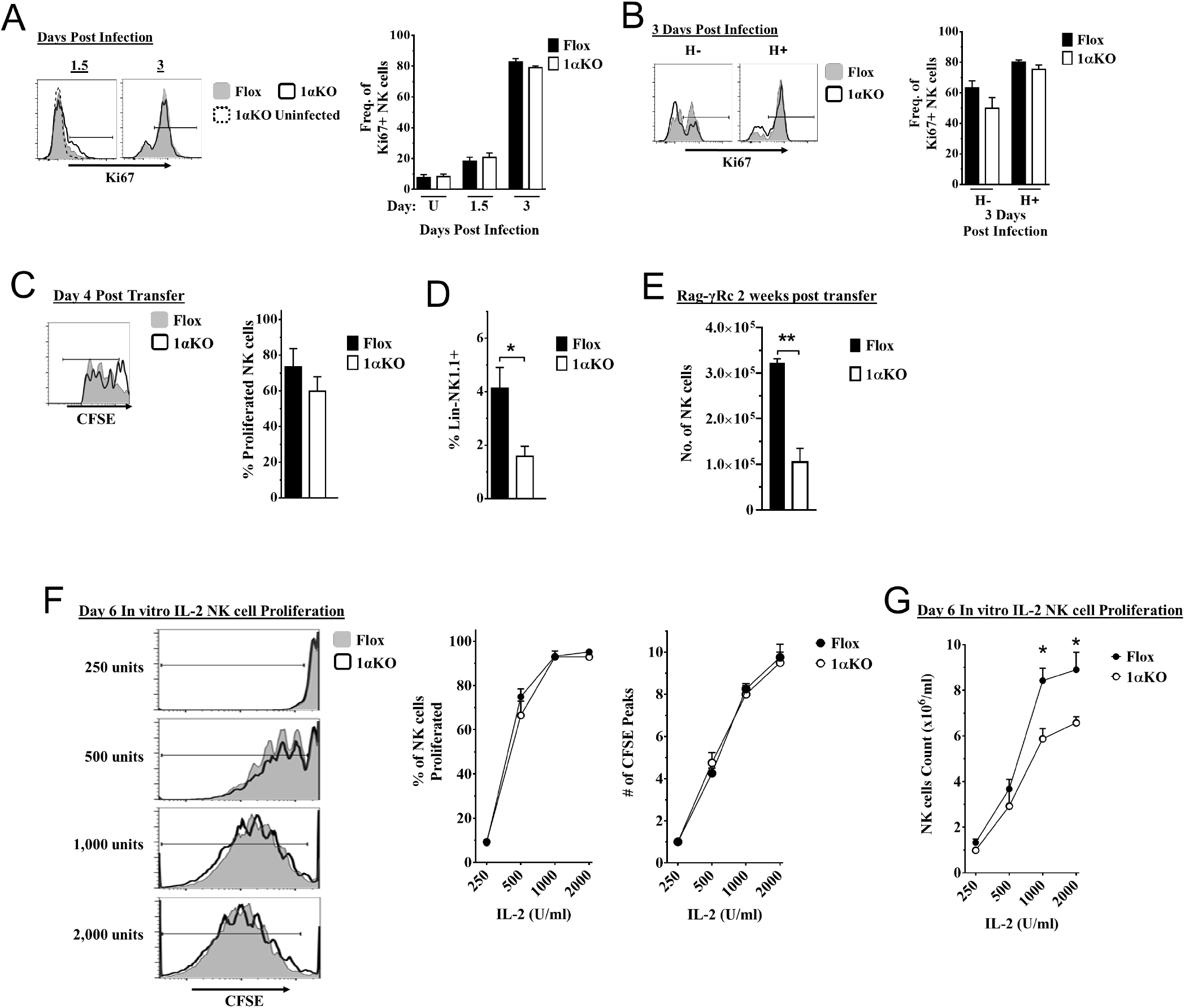
Cell division in HIF1αKO NK cells is normal but numbers are reduced. (A-B) 1αKO or FL control mice infected with MCMV received a dose of 50K p.f.u. and analyzed as indicated. (A) Ki67 expression with representative histograms (left) and frequency quantification (right) in 1αKO or FL control bulk NK cells at day 1.5 and day 3 pi. Data are from 3 independent experiments with 5-7 mice per group. (B) Expression of Ki67 in Ly49H negative and positive NK cells shown in histogram (left) and frequencies quantified (right). Data are from 2 independent experiments with 3 mice per group. (C&D) Rag-γRc mice analyzed for CFSE+ splenic NK cells from 1αKO or FL control mice 4 days post transfer, showing representative histogram (C, left) and quantification of total proliferation (C, right), and frequencies (D). Data are from 2 independent experiments with 4 mice per group. (E) Rag-γRc mice analyzed on day 14 since post splenocyte transfer for numbers of NK cells in the spleen from 1αKO or FL control mice. Data are from 2 independent experiments with 3 mice per group. (F-G) In vitro proliferation assays were done using human recombinant IL-2 at different concentrations and time points indicated below. (F) Representative histograms (left), quantification of total proliferated cells (middle), and number of CFSE peaks (right) of CFSE labeled NK cells from 1αKO or FL control mice stimulated with different concentrations of IL-2 for 6 days. (G) Numbers of NK cells were counted from figure F and graphed in cells per one milliliter. Data are from 2 independent experiments with 4 mice per group. Data depict mean +/− SEM, with each data set containing data indicated number of mice per group from independent experiments. Unpaired t-test was performed on A-G. Statistical significance indicated by n.s., no significant difference; * = *p*<.05; ** = *p*<.01; *** = *p*<.001; **** = *p*<.0001.

At later time points, Ly49H^+^ NK cells demonstrate selective proliferation, resulting in expansion [6, 7]. Since Ly49H expansion was reduced in 1α KO mice (**Figure 2D**), we specifically assessed proliferation of Ly49H^+^ NK cells. Surprisingly, Ki67 expression on Ly49H^+^ cells in 1αKO mice was comparable to FL controls (**Figure 4B**). Furthermore, the frequencies of Ly49H^+^ NK cells in both groups of mice were comparable at day 3 pi (**Figure S2B**). At the peak of Ly49H^+^ NK cell expansion at day 7 pi, there were again no differences in frequencies of Ki67 expression in 1αKO and FL controls (**Figure S2C, S2D**). Thus, during MCMV infection, selective Ly49H^+^ NK cell proliferation is also normal in NK cells lacking HIF1α.

To extend our analysis of NK cell proliferation *in vivo* in the absence of HIF1α, we utilized a non-inflammatory *in vivo* model for homeostatic NK cell proliferation by adoptively transferring 1αKO or FL splenocytes into lymphopenic Rag-γRc mice. CFSE-labeled, HIF1α-sufficient and -deficient splenic NK cells analyzed at day 4 post transfer showed no significant differences in cell division but the frequency of 1αKO NK cells was reduced compared to FL NK cells (**Figure 4E**). At two weeks post transfer, reduced numbers of 1αKO NK cells were again observed as compared to FL controls (**Figure 4F**) while all other immune cells showed comparable or even increased engraftment when derived from 1αKO mice (**Figure S2E**). Thus, HIF1α is dispensable for homeostatic proliferation in NK cells *in vivo* though ultimate expansion was affected.

Since cytokines such as IL-2 can induce NK cell proliferation and expansion, we tested their *in vitro* effects on HIF1α-deficient NK cells. Indeed, when enriched CFSE-labeled splenic 1αKO and FL NK cells were *in vitro* stimulated with different concentrations of IL-2, we found comparable cell division (**Figure 4F**). Interestingly, however, when NK cells were quantified after 6 days of culture, 1αKO NK cell numbers were significantly reduced compared to FL NK cells, though these differences were less evident at lower IL-2 concentrations (**Figure 4G**). Taken together, these data indicate that HIF1α controls NK cell numbers after proliferation though proliferation itself is HIF1α-independent.

### HIF1α KO NK cells are predisposed to apoptosis

Since 1αKO NK cells showed normal proliferation but did not expand, we hypothesized that HIF1α plays an unexpected role in NK cell survival. Since survival is mediated through signaling and interactions of pro-survival (Mcl-1, Bcl2) and pro-apoptotic proteins (Bim, Cytochrome c), we examined if these factors were associated with HIF1α. At 3 days after stimulation, IL-2 activated 1αKO NK cells showed significant increases in transcripts for pro-apoptotic Bim, cytochrome c, and caspase-3, as compared to FL controls (**Figure 5A**) which was coupled with reduced numbers (**Figure S3A**). Furthermore, Bim, and cytochrome c protein expression was significantly increased while protein expression of pro-survival Bcl-2 showed a significant decrease (**Figure 5B**). However, expression of a known HIF1α pro-survival gene target, Mcl-1[20-22], was not increased in 1αKO NK cells (**Figure S3C**). Thus, HIF1α plays an unexpected role in survival gene expression rather than proliferation in IL-2-stimulated NK cells *in vitro* suggesting this effect might account for decreased accumulation of Ly49H^+^ NK cells during MCMV infection *in vivo*, despite normal proliferation (**Figs 2D, 4A, 4B**).

**Figure 5.**
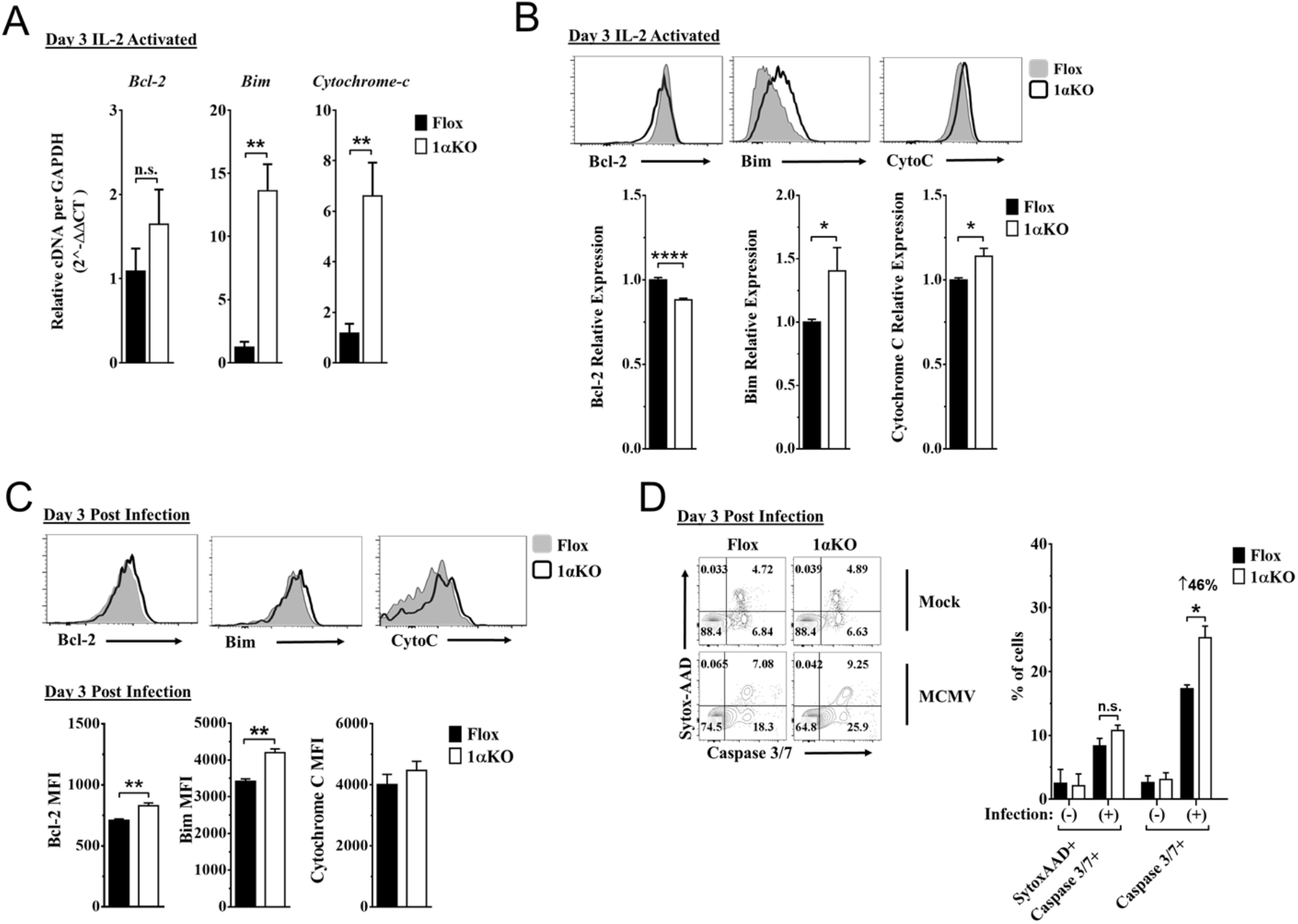
HIF1α KO NK cells are predisposed to apoptosis. (A) mRNA transcripts for levels of apoptosis associated genes Bcl-2, Bim, and Cytochrome c was measured in 1αKO or FL control NK cells at day three post IL-2 stimulation. Data are from 2 independent experiments with 4 mice per group. (B) Representative histograms (top) and quantification (bottom) of pro-survival relative protein expression of Bcl-2, Bim, and Cytochrome c (Cyto-C) in NK cells three days post IL-2 stimulation. Data are from 2-5 independent experiments with 4-11 mice per group. (C) Representative histograms (top) and quantification (bottom) of Bcl-2 MFI, Bim MFI, and Cytochrome c (Cyto-C) MFI in NK cells day 3 post MCMV infection. Data are from 2-3 independent experiments with 4-6 mice per group. (D) Representative contour plots (left) and frequency quantification (right) of Necrotic cells (Caspase 3/7+Sytox AAD+), and Apoptotic cells (Caspase 3/7+Sytox AAD-). Data are from 2 independent experiments with 3 mice per group. Data depict mean +/− SEM, with each data set containing data indicated number of mice per group from independent experiments. Unpaired t-test was performed on A-D. Statistical significance indicated by n.s., no significant difference; * = *p*<.05; ** = *p*<.01; *** = *p*<.001; **** = *p*<.0001.

Indeed, during MCMV infection *in vivo*, 1αKO NK cells showed elevated Bim expression at day 1.5 and day 3 pi (**Figure S3D and Figure 5C**, respectively), consistent with when HIF1a expression was highest (day 3 pi, **Figure 1F**) and the number of NK cells began to significantly decline (**Figure 2C,D**). We next sought to determine if there was increased cell death by staining for Caspase 3/7 and Sytox-AAD for apoptotic cells (single positive Caspase 3/7) and necrotic cells (double positive). At day 3 pi, 1αKO NK cells displayed increased apoptotic activity (46% increase compared to FL controls) when examined directly *ex vivo* with no difference in necrotic cells (**Figure 5D**). HIF1α regulates the autophagy gene BNIP3, raising the possibility that abnormal autophagy could account for our findings. However, no significant differences in the autophagy markers p62 and CytoID were detected between 1αKO and FL control NK cells at day 3 pi (**Figure S3E**). Taken together, these data show that HIF1α protein plays an important role in regulating survival, not proliferation, of NK cells during MCMV infection.

### Impaired glycolytic activity contributes to impaired fitness in HIF1α KO NK cells

Our data indicate that 1αKO NK cells do not have impaired cell division but rather show a predisposition for apoptosis that results in reduced numbers compared to FL controls during cytokine activation. We verified that HIF1α-deficient NK cells manifest increased oxidative phosphorylation (OXPHOS) (**Figure S3F**) that was previously reported but whose basis was not further evaluated [18]. Since these findings suggest that NK cell metabolism may be dysregulated in the absence of HIF1α, we ascertained if HIF1α-associated glycolytic genes were involved despite being considered dispensable [19, 23]. First, we determined if these metabolic changes could be due to glucose uptake defects since HIF1α affects glucose transport genes [24, 25]. However, despite verification of absence of HIF1α in 1αKO NK cells by direct measurement of HIF1α transcripts at 3 days post IL-2 stimulation (**Figure 6A**), we found that glucose uptake was normal as uptake of the 2-NDBG fluorescent glucose analog showed no defect (**Figure 6B**). By contrast, the expression of the HIF1α glycolytic target genes, hypoxanthine-guanine phosphoribosyl transferase (*Hrpt)*, glucose transporter type 1 (*Glut1*), pyruvate kinase M2 (*Pkm2*), hexokinase 2 (*Hk2*), glucose-6-phosphate isomerase 1 (*Gpi1*), lactate dehydrogenase A (*Ldha*), and phosphoinositide-dependent kinase 1 (*Pdk1*), were markedly reduced in 1α KO NK cells (**Figure 6C**). Furthermore, we found an impaired utilization of glucose and production of lactate when we measured glucose metabolism using liquid chromatography mass spectrometry (LCMS) (**Figure 6D**). However, no defects were found in glycolysis-independent metabolism, i.e., glutamine/glutamate pathway. Thus, HIF1α affects metabolism of NK cells by supporting glycolysis.

**Figure 6.**
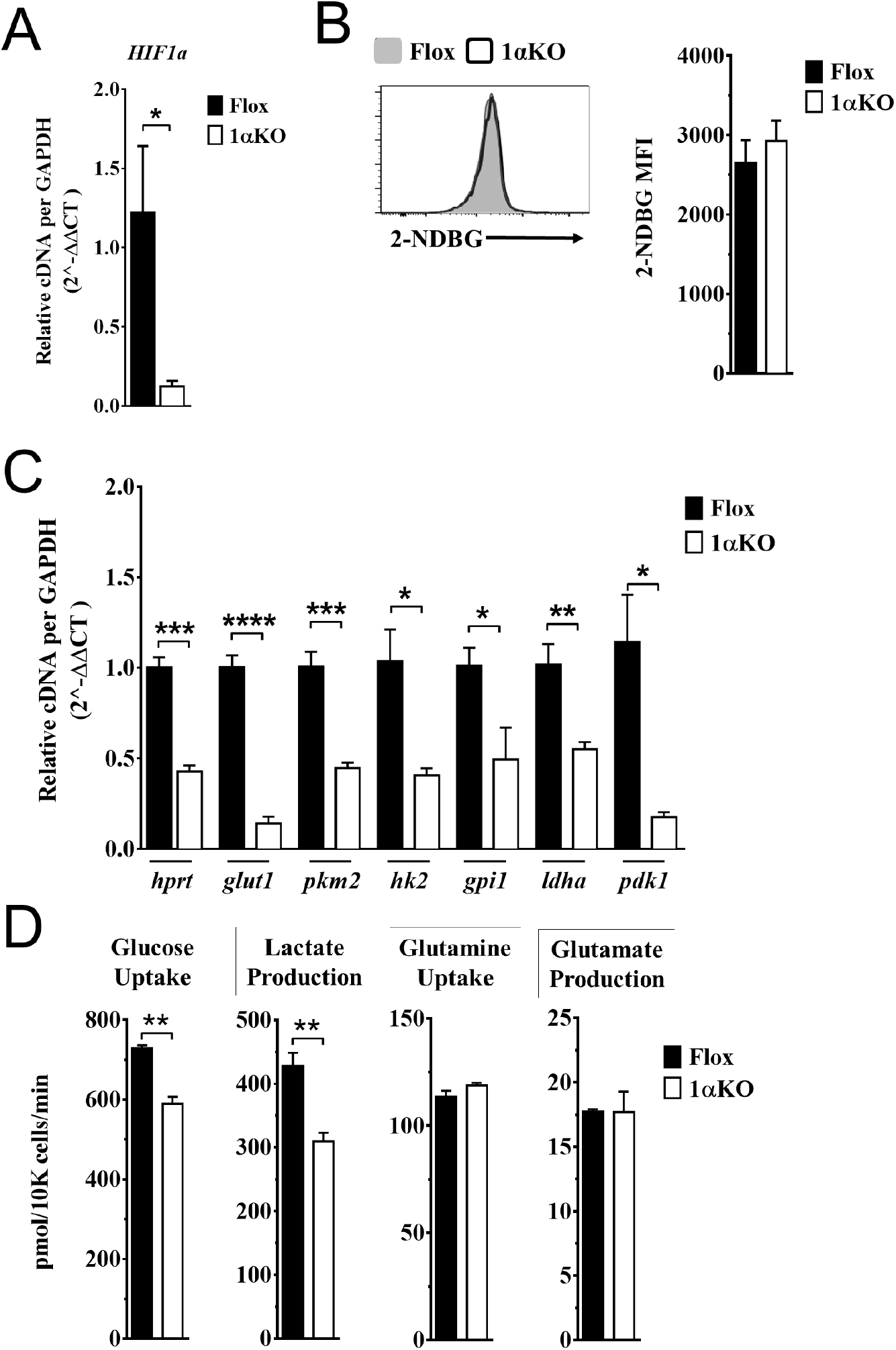
Impaired glycolytic activity contributes to impaired survival of HIF1α KO NK cells. Enriched NK cells from 1αKO or FL control mice were IL-2 treated for 3 days then, collected, and analyzed as follows:(A) mRNA transcript levels of HIF1α for both cohort of mice measured with qRT-PCR. Data are from 2 independent experiments with 4 mice per group. (B) Flow cytometry analysis of 2-NDBG MFI expression as a measure of glucose intake. Data are from 3 independent experiments with 6 mice per group. (C) mRNA transcripts levels for glycolysis genes glut1, gpi1, hk2, hprt, ldha, pdk1, and pkm2 were measured with qRT-PCR. Data are from 2 independent experiments with 4 mice per group. (D) Media was collected from day 3 IL-2 activated NK cell cultures and using LC/MS, uptake of glucose (1^st^ panel) and glutamine (3^rd^ panel), and production of lactate (2^nd^ panel) and glutamate (4^th^ panel) was measured. Data are representative of 2 independent experiments with 3 mice per group. Data depict mean +/− SEM, with each data set containing data indicated number of mice per group from independent experiments. Unpaired t-test was performed on A-D. Statistical significance indicated by n.s., no significant difference; * = *p*<.05; ** = *p*<.01; *** = *p*<.001; **** = *p*<.0001.

## Discussion

Here we define a previously undescribed role for HIF1α in supporting NK cell-mediated protection against MCMV infection. In the selective absence of HIF1α in NK cells, we found increased morbidity and viral burden. However, we found HIF1α KO NK cells displayed normal activation and proliferation in response to MCMV but they did not normally expand in number. This defect was evident in another *in vivo* response, i.e., homeostatic proliferation in lymphopenic mice, as well as with *in vitro* IL2 stimulation where there was normal cell division but impaired expansion. The 1αKO NK cells had an increase in apoptosis as a result of decreased expression of glycolytic genes and abnormal glucose metabolism that have been previously associated with apoptosis [12, 26-28]. Thus, HIF1α supports survival in NK cells through providing optimal glucose metabolism during pathogen infection.

While HIF1α protein function in NK cells during pathogen infection had not been detailed previously, it was recently explored in tumor-infiltrating NK cells in two studies with comparable results attributed to different mechanisms [17, 18]. In mice with HIF1α-deficient NK cells, tumor control was enhanced either directly by increased IL-18 signaling that enhanced NK cell effector responses, or indirectly through an absence of tumor-infiltrating NK cells which altered the VEGF/VEGFR axis affecting productive angiogenesis. In contrast, our findings were the polar opposite to these reports in that mice with HIF1α-deficient NK cells showed reduced control of viral infection without alteration in NK cell activation or killing ability. While HIF1α-deficient NK cell numbers were normal in the naïve spleen, after MCMV infection there was a significant decrease in numbers with impaired expansion of Ly49H^+^ cells that we attribute to apoptosis. Therefore, the function of HIF1α in NK cells markedly differs in pathogen responses, as compared to NK cell responses to tumors.

Here we provide evidence that the absence of HIF1α affects survival in NK cells during a pathogen infection. HIF1α senses intracellular changes and supports transcription of genes related to survival in several cellular pathways [12, 29-31]. In the apoptosis pathway, constitutively expressed pro- and anti-apoptotic proteins antagonize each other to sustain viability in normal healthy cells and when these proteins are unbalanced due to external or internal stimuli, pro-apoptotic proteins are preferentially activated and expressed, resulting in the initiation of programmed cell death via caspase activity [26, 32, 33]. In tumor models, HIF1α can directly impact apoptosis by upregulating the pro-survival gene myeloid cell leukemia sequence-1 (Mcl-1) [20-22]. Mcl-1 was found to be highly expressed in normal NK cells and genetic ablation of Mcl-1 in NK cells resulted in a dramatic loss of mature NK cells at baseline that spared immature populations [34]. This absence of NK cells impaired control of tumors as well as protection against lethal sepsis. Here Mcl-1 is unlikely to contribute to the HIF1α-deficient NK cell phenotype because we found no difference in transcript or protein levels of Mcl-1 in 1αKO compared to FL control NK cells, and there were no baseline differences in NK cell numbers. Another potential survival pathway regulated by HIF1α in tumor cells is through the increased expression of BCL2/adenovirus E1B 19 kDa protein-interacting protein 3 (BNIP3), a component of autophagosomes that complexes with p62 LC3 complex proteins to initiate a survival mechanism of self-renewal. In MCMV infection, O’Sullivan et al. reported that BNIP3 supported memory formation in NK cells [35]. Wild-type (WT) Ly49H+ NK cells were transferred into Ly49H-deficient mice; upon infection, memory NK cells displayed levels of BNIP3 transcripts comparable to resting NK cells. However, in a competition experiment, *Bnip3*^−/−^ NK cells showed impaired survival during the contraction phase as compared to WT NK cells. The markers CytoID and LC3 that indicated active autophagy were comparable between resting and memory WT NK cells but autophagy was impaired in memory *Bnip3*^−/−^ NK cells which displayed higher CytoID expression. Interestingly, BNIP3 transcripts were reduced during the effector phase of the response but this was not explored and the interplay between BNIP3 and HIF1α was not determined. In our studies, there were no differences in CytoID and p62 at three days post MCMV infection in 1α KO and FL NK cells, suggesting this pro-survival axis is unaffected in the absence of HIF1α. Finally, when survival proteins are unaltered, a mechanism of subverting survival is the over-expression of pro-apoptotic proteins. Multiple pathways converge on the potent inducer of apoptosis, Bim, which stimulates caspase activity [26, 32, 33]. Bim regulates NK cell survival and restrains the expansion of memory cells during MCMV infection [36, 37]. Bim can potentiate apoptosis even in the presence of sufficient pro-survival signals in NK cells but the interplay between HIF1α and Bim was not previously explored [38]. In our experiments, HIF1α-deficient NK cells had elevated levels of Bim as compared to controls during *in vitro* proliferation and MCMV infection. Furthermore, HIF1α-deficient NK cells showed a marked increase in caspase activity, a direct measurement of apoptosis, as compared to Flox controls at three days post infection. Thus, in our studies we found activated HIF1α-deficient NK cells upregulate the key pro-apoptotic protein Bim and its downstream targets during *in vitro* proliferative and pathogen infection responses to promote cell death while there was little evidence implicating other survival pathways.

Our studies indicate HIF1α-deficient NK cells have impaired metabolism of glucose that provides fuel for glycolysis, critical for cellular survival [12, 26-28, 31, 39]. The importance of glycolysis in NK cells has been previously studied using glycolytic inhibitors. Keppel et al. and Mah et al. inhibited glucose metabolism during acute activation, and during MCMV infection that resulted in a reduction of IFNγ responses, and impaired proliferation and protective immunity, respectively [40, 41]. However, data from our studies here and others [18, 42] suggest HIF1α-associated glucose metabolism is dispensable for effective IFNγ responses in NK cells. Moreover, Loftus et al reported that HIF1α was not required for NK cell activation, transcription of glycolytic genes, and glucose metabolism [19]. Further, it was determined that cMyc was the potent regulator of glucose metabolism, leading to the conclusion that HIF1α was dispensable in NK cells for glucose metabolism [23]. One potential important caveat for these studies is that NK cells were expanded for six days in IL-15 *in vitro* then stimulated with IL-2 and IL-12 for activation [19]. In another *in vitro* study, enriched HIF1α-deficient NK cells were IL-2 stimulated for seven days in hypoxic conditions then stimulated for 4-6 hours with IL-12 and IL-18 [18]; deletion of HIF1α in NK cells resulted in higher OXPHOS, presumably to compensate impaired glucose metabolism. However, in this same experiment, glucose metabolism was comparable in both HIF1α-deficient and - sufficient NK cells. In our studies here, we measured metabolism directly *ex vivo* and found increases in both glucose metabolism and OXPHOS in resting HIF1αKO NK cells compared to Flox controls. Thus, prior results were skewed by culture conditions that likely do not reliably represent *in vivo* effects of HIF1α on glucose metabolism in NK cells.

In further contrast to aforementioned studies, we found that in the absence of HIF1α in NK cells, IL-2 stimulation resulted in a reduced glycolytic transcriptome and a subsequent reduction in glucose metabolism. These experiments coincide with other IL-2 stimulation assays we performed where pro-apoptotic Bim was upregulated at the gene and protein level. Importantly, reduced glucose metabolism has been directly associated with apoptosis via Bim-caspase-3 pathway [26, 28, 32, 43]. In summary, expression of HIF1α gene targets is necessary to provide sufficient glucose metabolism to support survival by repressing Bim protein expression thereby sustaining adequate numbers of NK cells for protective immunity.

Targeting HIF1α using small molecule inhibitors has drawn considerable interest over recent years for treatment of multiple pathologies. However, HIF1α is ubiquitously expressed and has pleiotropic properties through targeting of multiple genes, as illustrated here and elsewhere [12, 13]. Therefore, special care should be taken when considering the use of HIF1α inhibitors as they may compromise NK cell function under certain conditions such as pathogen infection.

## Materials and Methods

### Mice

Wild type B6 mice (C57/BLNCR1) were purchased through Charles River Laboratories (Wilmington, MA). HIF1α specific deletion in NK cells was achieved through crossing B6.129-*Hif1a*^*tm3Rsjo*^/J mice from The Jackson Laboratory (Bar Harbor, ME) with *Ncr1*^iCre^ mice from Eric Vivier. Control β2m knock out mice were obtained from The Jackson Laboratory.

### Splenocyte and NK Cell Culture

Spleens from mice were collected and placed in complete RPMI (Gibco#11875 RPMI 1640, 10% fetal calf serum, 1% penicillin/streptomycin) and minced through 100μm filter to create single cell suspensions. Splenocytes were then RBC lysed with lysis buffer (1M NH_4_NaCl, 1M HEPES) for 2 minutes and washed with complete RPMI. Splenocytes were counted and cultured at 10×10^6^ cells per ml in complete RPMI, 10μM non-essential amino acids, 0.57μM 2-mercaptoethenol. When NK cell enrichment was used, NK cells were isolated from splenocytes using Stem Cell Technologies NK isolation kit (#19855) as described by manufacturer. Cytokines were obtained from Peprotech: IL-15 (210-15), IL-18 (B002-5), Pestka Biomedical Laboratories assay science IFNβ (12401-1) and used as indicated in figure legends. Splenocyte cultures were incubated at 37°C in a normal incubator (20% O_2_) or hypoxic chamber (1% O_2_). For intracellular cytokine measurements, NK cells were cultured for 20 hours then brefeldin A was added for the last four hours.

### Nutrient Uptake Analysis

For nutrient uptake analysis with LC/MS, enriched NK cells from FL and 1α KO mice were cultured in 96-well plate with media containing 2000 units of recombinant human IL-2. Media were refreshed on day 1. On day 3 after a further 48 hr-incubation, the spent media were collected and analyzed. Known concentrations of U-^13^C internal standards (glucose, lactate, glutamine, and glutamate; Cambridge Isotopes) were spiked into media samples before extraction. Extractions were performed in glass to avoid plastic contamination as previously reported [44]. Media samples were measured by liquid chromatography coupled mass spectrometry (LC/MS) analysis, with a method previously described [45]. Samples were run on a Luna aminopropyl column (3 μm, 100 mm × 2.0 mm I.D., Phenomenex) in negative mode with the following buffers and linear gradient: A = 95% water, 5% acetonitrile (ACN), 10 mM ammonium hydroxide, 10 mM ammonium acetate; B = 95% ACN, 5% water; 100% to 0% B from 0-45 min and 0% B from 45-50 min; flow rate 200 μL/min. Mass spectrometry detection was carried out on a Thermo Scientific Q Exactive Plus coupled with ESI source. For each compound, the absolute concentrations were determined by calculating the ratio between the fully unlabeled peak from samples and the fully labeled peak from standards. The consumption rates were normalized by cell growth over the experimental time period [46].

### Seahorse Studies

NK cells were enriched from FL and 1α KO mice and 4×10^5^ cells were transferred into 96 well plate without any stimulation. Agilent Seahorse XF Cell Mito Stress test was performed on NK cells using oligomycin (10 μM), FCCP (10 μM), rotenone (10 μM) plus antimycin A (10 μM) over two hours. Oxygen consumption rate (OCR) and extracelluar acidification rate (ECAR) was measured analyzed using Agilent Seahorse XFe96 Analyzers.

### MCMV infection

MCMV stocks were derived from salivary gland homogenates and titered using 3T12 (ATCC CCL-164) fibroblast cell line. The complete genomic sequence for the strain used here was previously published by our lab [47]. Virus was inoculated intraperitoneally (IP) in a total volume of 200μl PBS at indicated plaque forming unit (pfu.) doses and mouse spleens were harvested at indicated time points for analysis. Morbidity was measured in weight loss everyday post infection for seven days. Viral load was measured as previously described [48].

### In vivo cytotoxicity assay

Target splenocytes were isolated from β2m knock out mice and labeled with 2μM CFSE (Life Technologies) and WT C57BL/6 controls were labeled with 10μM CFSE. CFSE^low^ and CFSE^high^ target cells were mixed at a 1:1 ratio and 2×10^6^ target cells were injected i.v. into FL or 1αKO mice that had received PBS or Poly:I-C inoculations three days prior. After 3 hours splenocytes were harvested and stained. The ratio of CFSE^low^ to CFSE^high^ viable bulk CD45^+^ splenocyte cells was determined by flow cytometry. Target cell rejection was calculated using the formula [(1− (Ratio (CFSE^low^: CFSE^high^) sample / Ratio (CFSE^low^: CFSE^high^) average of NK depleted)) × 100].

### Flow cytometry

After preparation, cells were stained in staining buffer (PBS, 0.2% Fetal Bovine Serum, 0.01% sodium azide), and NK cells were identified as e780-CD45+CD3-CD19-NK1.1+NKp46+. Antibodies and reagents from eBioscience: anti-CD3ε (145-2C11), anti-CD19 (eBio1D3), anti-NK1.1 (PK136), anti-NKp46 (29A1.4), anti-CD11b (M1/70), anti–IFN-γ (XMG1.2), anti-KLRG1(2F1), anti-NKG2A.B6 (16a11), anti-Ly49G2 (eBio4D11), anti-Ly49H (3D10), anti-Ly49I (YLI-90), anti-CD69 (H1.2F3), Fixable Viability Dye eFluor 780 (65-0865-14), anti-Ki67 (48-5698-82), Brefeldin A (00-4506-51), Transcription Factor Staining kit (00-5521), Invitrogen: Granzyme B (MHGB4); BD Pharmingen: anti-Ly49A (116805), BrdU (51-23614L), BD Biosciences: BD Cytofix/Cytoperm kit (554714), Biolegend: anti-NK1.1 (PK136), anti-HIF1α (546-16), anti-Bcl-2 (10C4), anti-cytochrome c (6H2,B4), anti-DNAM-1 (132006); Jackson ImmunoResearch: Alexa Fluor 647 F (ab’)_2_ Fragment Goat Anti-Rabbit IgG (111-606-046), RC fragment specific; Invitrogen: anti-Bim (K.912.7), CYTO-ID (ENZ-51031-0050), Cell Signaling: SQSTM1/p62 antibody (5114-5114S), Mcl-1 (65617S). For caspase activity assay, splenic NK cells were identified being CD45^+^, Lineage negative (CD3^−^ CD19^−^) NKp46^+^ and stained as indicated by manufacturer’s protocol with Cell Event Caspase 3/7 Green Flow Cytometry Assay Kit (Molecular Probes). All samples were analyzed using BD Canto FACS instrument ((BD Biosciences) Flow cytometry files were analyzed using FlowJo v10 software (Tree Star).

### Analysis of RNA

RNA levels from enriched NK cells were measured using quantitative reverse transcriptase PCR (qRT-PCR). RNA was isolated from cells using Qiagen RNeasy Plus Mini Kit (Qiagen). cDNA was synthesized using ∼1μg of DNase-treated RNA with random hexamers and SuperScript III reverse transcriptase (Invitrogen). qPCR of cDNA was performed using PowerSYBR Green PCR Master Mix (Applied Biosystems) using the following primers: *Bcl-2* (5’-CCTGTGGATGACTGAGTACCTG-3’and 5’-AGCCAGGAGAAATCAAACAGAGG-3’), *Bim* (5’-GGAGATACGGATTGCACAGGAG-3’and 5’- CTCCATACCAGACGGAAGATAAAG-3’), *Cytochrome-c* (5’-GAGGCAAGCATAAGACTGGACC-3’and 5’-ACTCCATCAGGGTATCCTCTCC-3’), *HIF1a* (5’-CATCAGTTGCCACTTCCCCA-3’and 5’-GGCATCCAGAAGTTTTCTCACAC-3’), *hprt* (5’-CTGGTGAAAAGGACCTCTCGAAG-3’ and 5’-CCAGTTTCACTAATGACACAAACG-3’), *glut1* (5’-AAGAAGCTGACGGGTCGCCTCATGC-3’and 5’-TGAGAGGGACCAGAGCGTGGTG-3’), *pkm2* (5’-CAGGAGTGCTCACCAAGTGG-3’and 5’-CATCAAGGTACAGGCACTACAC-3’), *hk2* (5’-GGAGAGCACGTGTGACGAC-3’and 5’-GATGCGACAGGCCACAGCA-3’), *gpi1* (5’-GTTGCCTGAAGAGGCCAGG-3’and 5’-GCTGTTGCTTGATGAAGCTGATC-3’), *ldha* (5’-CACAAGCAGGTGGTGGACAG-3’and 5’-AACTGCAGCTCCTTCTGGATTC-3’), *pdk1* (5’-GATTCAGGTTCACGTCACGCT-3’and 5’-GACGGATTCTGTCGACAGAG-3’), *Mcl-1* (5’-AGCTTCATCGAACCATTAGCAGAA-3’ and 5’-CCTTCTAGGTCCTGTACGTGGA-3’), and *GAPDH* (5’-GTTGTCTCCTGCGACTTCA-3’and 5’-GGTGGTCCAGGGTTTCTTA-3’). qPCR was performed using StepOnePlus Real-Time PCR System (Applied Biosystems). Transcript abundance relative to GAPDH was calculated using delta-delta CT.

### Statistics

Graphed data are represented as means with standard error of the mean (SEM). Statistics were determined using Graphpad software, and 1-way ANOVA was used when comparing multiple groups or unpaired student *t*-test when comparing two groups as indicated in the figure legends. Statistical significance indicated by n.s., no significant difference; * = *p*<.05; ** = *p*<.01; *** = *p*<.001; **** = *p*<.0001.

---

**Supplement 1.**
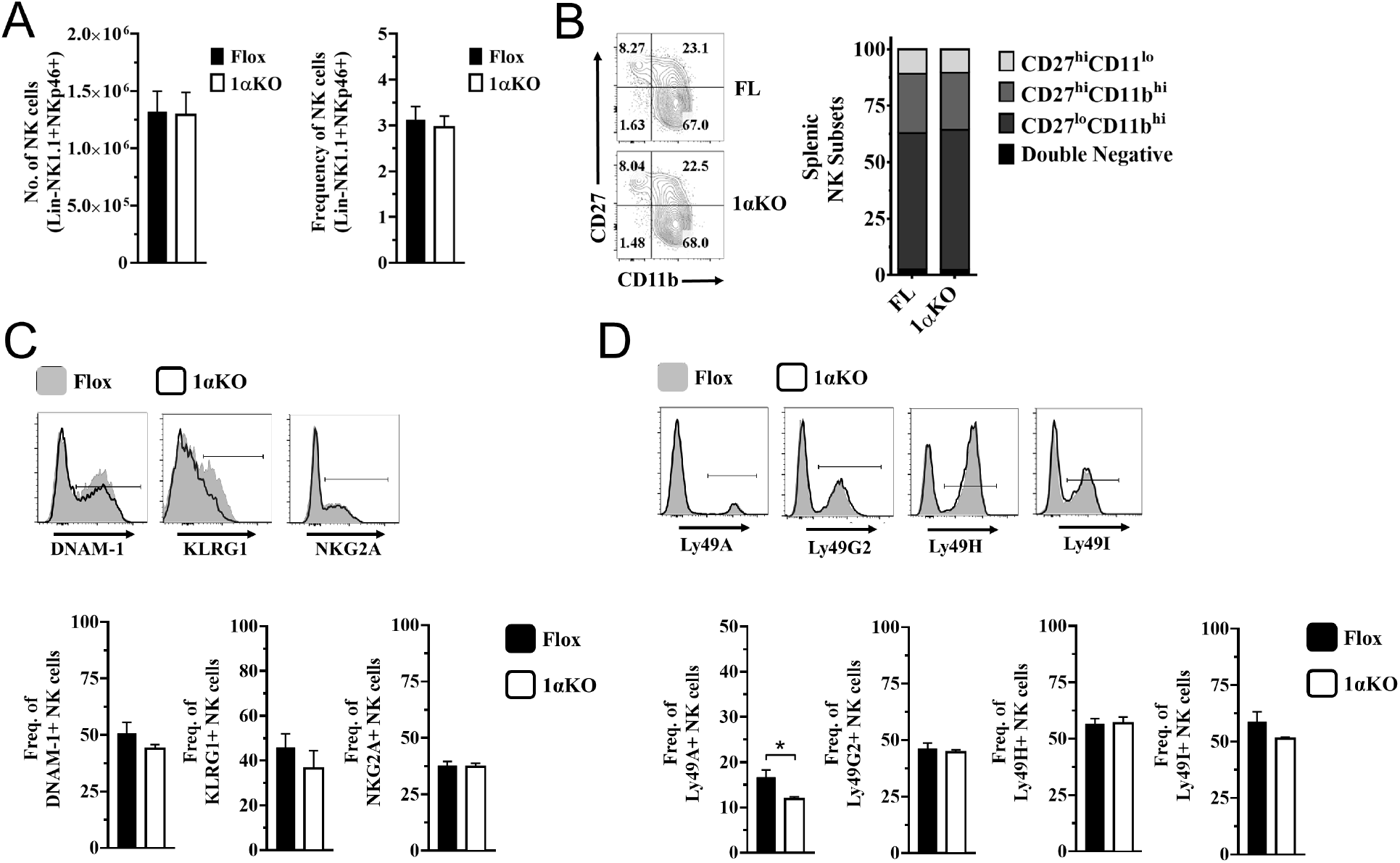
HIF1α expression in NK cell and effect of genetic deletion. (A) Quantification of NK cells absolute numbers (left) and frequencies (right) in 1αKO or FL control mice direct *ex vivo*. Data are from 3 independent experiments with 6-9 mice per group. (B) Representative contour plots (left) and quantification (right) of CD27 and CD11b maturation markers direct *ex vivo* in NK cells of 1αKO or FL control mice. Data are from 2 independent experiments with 3 mice per group. (C) Representative histograms (top) and frequencies (bottom) of DNAM-1, KLRG1, and NKG2A on NK cells of 1αKO or FL control mice. Data are from 2-3 independent experiments with 3-7 mice per group. (D) Representative histograms (top) and frequencies (bottom) of Ly49A, Ly49G2, and Ly49H receptors on NK cells of 1αKO or FL control mice. Data are from 2-3 independent experiments with 3-7 mice per group. Data depict mean +/− SEM, with each data set containing data indicated number of mice per group from independent experiments. Unpaired t-test was performed on A-C. Statistical significance indicated by n.s., no significant difference; * = *p*<.05; ** = *p*<.01; *** = *p*<.001; **** = *p*<.0001.

**Supplementary 2.**
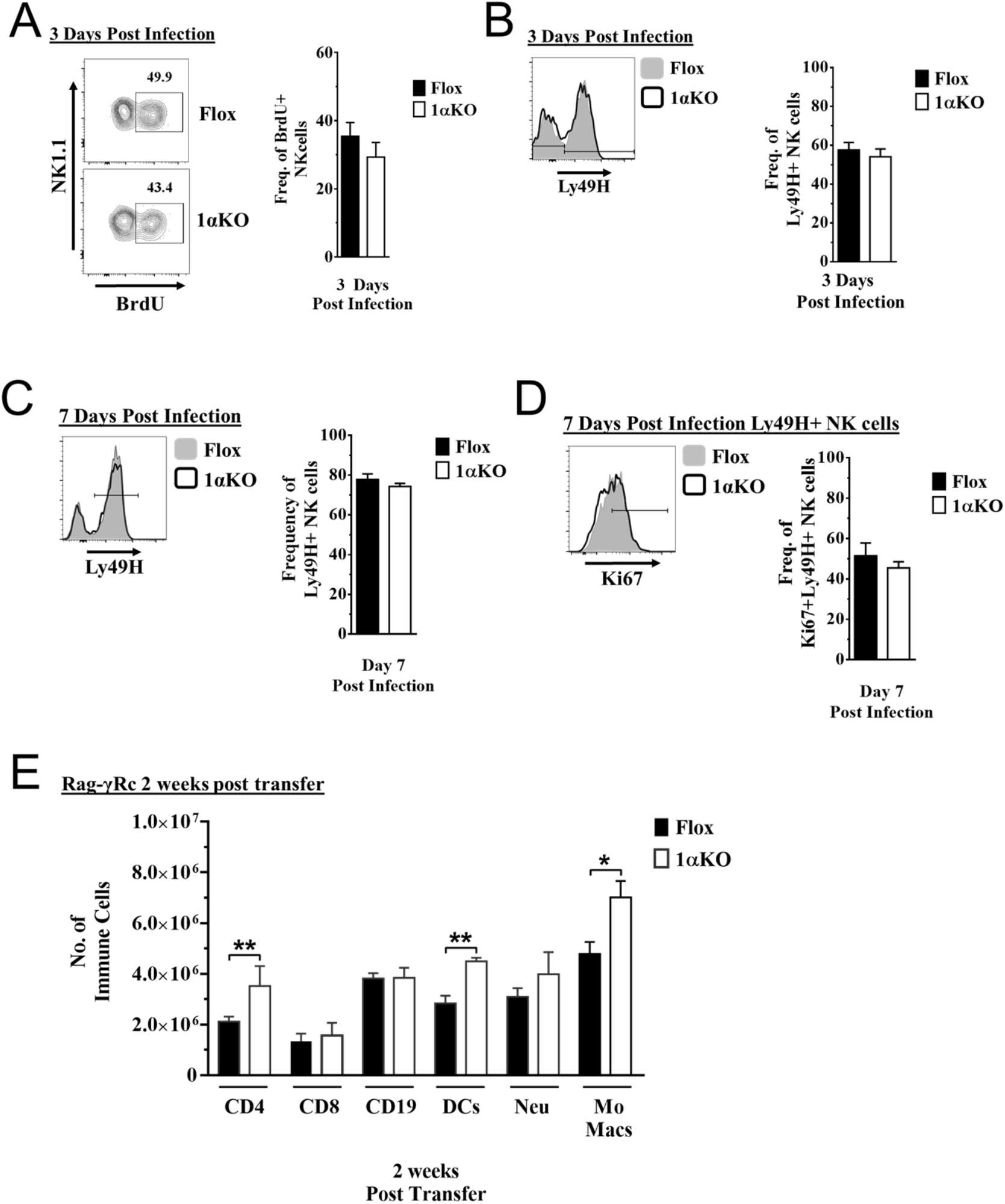
HIF1αKO NK cells proliferate comparable to Flox controls but have reduced numbers. (A-D) 1αKO or FL control mice infected with MCMV received a dose of 50K p.f.u. and analyzed as indicated. (A) Representative contour plots (left) and quantification (right) of bulk splenic NK cells that were collected day 3 post MCMV infection for BrdU expression. Data are from 2 independent experiments with 4 mice per group. (B) Representative histogram (left) and quantification of frequency (right) of Ly49H^+^ NK cells at day 3 post MCMV infection. Data are from 2 independent experiments with 4 mice per group. (C) Representative histogram (left) and quantification of frequency (right) of Ly49H expressing NK cells at day 7 post MCMV infection. Data are from 4 independent experiments with 7 mice per group. (D) Histogram that is representative of Ki67 expression in Ly49H^+^ NK cells and quantification at day 7 post MCMV infection. Data are from 4 independent experiments with 7 mice per group. (E) Rag-γRc mice analyzed on day 14 since post splenocyte transfer for engraftment of lymphocytes and myeloid cells in the spleen. Data are from 2 independent experiments with 3 mice per group. Data depict mean +/− SEM, with each data set containing data indicated number of mice per group from independent experiments. Unpaired t-test was performed on A-E. Statistical significance indicated by n.s., no significant difference; * = *p*<.05; ** = *p*<.01; *** = *p*<.001; **** = *p*<.0001.

**Supplementary 3.**
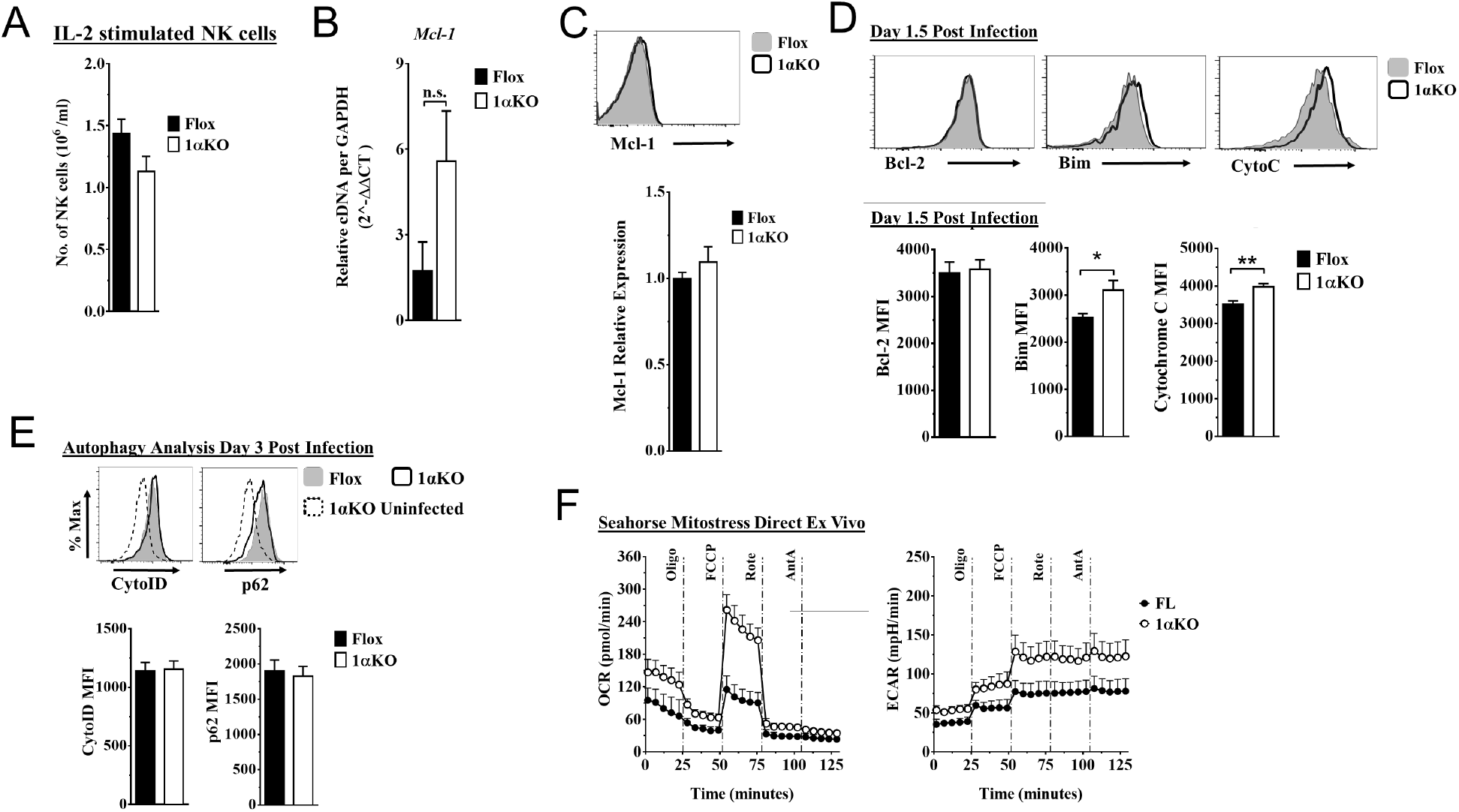
HIF1α KO NK cells are predisposed to apoptosis. (A-C) Enriched NK cells from 1αKO or FL control mice were stimulated with 2000 units of IL-2 for three days were then harvested and analyzed as indicated. Data represent 4 mice per time point from 2 independent experiments. (A) Quantification of the number of NK cells in culture at day 3. Data are from 2 independent experiments with 4 mice per group. (B) mRNA levels for Mcl-1 were quantified using qRT-PCR. Data are from 2 independent experiments with 4 mice per group. (C) Representative histogram (top) and quantification of relative protein expression pro-survival marker Mcl-1. Data are from 2 independent experiments with 4 mice per group. (D) Representative histograms (top) and quantification (bottom) of pro-survival protein Bcl-2 MFI, and pro-apoptotic proteins Bim MFI, and Cytochrome c (Cyto-C) MFI in NK cells at day 1.5 post MCMV infection. Data are from 2-3 independent experiments with 4-6 mice per group. (E) Autophagy markers CytoID and p62 were measured day 3 post MCMV infection. Data are from 2 independent experiments with 4-5 mice per group. (F) Enriched NK cells from 1αKO or FL control mice were subjected to a mitochondrial stress test and oxygen consumption rate (OCR) and extracellular acidification rate (ECAR) were measured direct ex vivo. Data are from 3 independent experiments with 5-6 mice per group. Data depict mean +/− SEM, with each data set containing data indicated number of mice per group from independent experiments. Unpaired t-test was performed on A-F. Statistical significance indicated by n.s., no significant difference; * = *p*<.05; ** = *p*<.01; *** = *p*<.001; **** = *p*<.0001.

## References

1. <SmithHR.YokoyamaWM 2002.PNAS Recognition of a virus-encoded ligand by a NK cell activation receptor MCMV.pdf>.

2. Arase, H., et al., Direct recognition of cytomegalovirus by activating and inhibitory NK cell receptors. Science, 2002. 296(5571): p. 1323–6.

3. Biron, C.A. and M.L. Tarrio, Immunoregulatory cytokine networks: 60 years of learning from murine cytomegalovirus. Med Microbiol Immunol, 2015. 204(3): p. 345–54.

4. Mitrovic, M., et al., Innate immunity regulates adaptive immune response: lessons learned from studying the interplay between NK and CD8+ T cells during MCMV infection. Med Microbiol Immunol, 2012. 201(4): p. 487–95.

5. Pyzik, M. and S.M. Vidal, NK cells stroll down the memory lane. Immunology & Cell Biology, 2009. 87(4): p. 261–263.

6. Dokun, A.O., et al., Specific and nonspecific NK cell activation during virus infection. Nat Immunol, 2001. 2(10): p. 951–6.

7. Sun, J.C., J.N. Beilke, and L.L. Lanier, Adaptive immune features of natural killer cells. Nature, 2009. 457(7229): p. 557–61.

8. Graham, A.M. and J.S. Presnell, Hypoxia Inducible Factor (HIF) transcription factor family expansion, diversification, divergence and selection in eukaryotes. PLoS One, 2017. 12(6): p. e0179545.

9. Kaelin, W.G., The von Hippel–Lindau Tumor Suppressor Protein. Annual Review of Cancer Biology, 2018. 2(1): p. 91–109.

10. Schofield, C.J. and P.J. Ratcliffe, Oxygen sensing by HIF hydroxylases. Nat Rev Mol Cell Biol, 2004. 5(5): p. 343–54.

11. Semenza, G.L., Oxygen sensing, hypoxia-inducible factors, and disease pathophysiology. Annu Rev Pathol, 2014. 9: p. 47–71.

12. Dengler, V.L., M. Galbraith, and J.M. Espinosa, Transcriptional regulation by hypoxia inducible factors. Crit Rev Biochem Mol Biol, 2014. 49(1): p. 1–15.

13. Schodel, J., et al., High-resolution genome-wide mapping of HIF-binding sites by ChIP-seq. Blood, 2011. 117(23): p. e207–17.

14. Baginska, J., et al., The critical role of the tumor microenvironment in shaping natural killer cell-mediated anti-tumor immunity. Front Immunol, 2013. 4: p. 490.

15. Balsamo, M., et al., Hypoxia downregulates the expression of activating receptors involved in NK-cell-mediated target cell killing without affecting ADCC. Eur J Immunol, 2013. 43(10): p. 2756–64.

16. Chambers, A.M. and S. Matosevic, Immunometabolic Dysfunction of Natural Killer Cells Mediated by the Hypoxia-CD73 Axis in Solid Tumors. Front Mol Biosci, 2019. 6: p. 60.

17. Krzywinska, E., et al., Loss of HIF-1alpha in natural killer cells inhibits tumour growth by stimulating non-productive angiogenesis. Nat Commun, 2017. 8(1): p. 1597.

18. Ni, J., et al., Single-Cell RNA Sequencing of Tumor-Infiltrating NK Cells Reveals that Inhibition of Transcription Factor HIF-1alpha Unleashes NK Cell Activity. Immunity, 2020. 52(6): p. 1075–1087 e8.

19. Loftus, R.M., et al., Amino acid-dependent cMyc expression is essential for NK cell metabolic and functional responses in mice. Nat Commun, 2018. 9(1): p. 2341.

20. Carrington, E.M., et al., Anti-apoptotic proteins BCL-2, MCL-1 and A1 summate collectively to maintain survival of immune cell populations both in vitro and in vivo. Cell Death Differ, 2017. 24(5): p. 878–888.

21. Liu, X.H., et al., HIF-1alpha has an anti-apoptotic effect in human airway epithelium that is mediated via Mcl-1 gene expression. J Cell Biochem, 2006. 97(4): p. 755–65.

22. Piret, J.P., et al., Hypoxia-inducible factor-1-dependent overexpression of myeloid cell factor-1 protects hypoxic cells against tert-butyl hydroperoxide-induced apoptosis. J Biol Chem, 2005. 280(10): p. 9336–44.

23. O’Brien, K.L. and D.K. Finlay, Immunometabolism and natural killer cell responses. Nat Rev Immunol, 2019. 19(5): p. 282–290.

24. Chen, C., et al., Regulation of glut1 mRNA by hypoxia-inducible factor-1. Interaction between H-ras and hypoxia. J Biol Chem, 2001. 276(12): p. 9519–25.

25. Huang, Y., et al., Normal glucose uptake in the brain and heart requires an endothelial cell-specific HIF-1alpha-dependent function. Proc Natl Acad Sci U S A, 2012. 109(43): p. 17478–83.

26. El Mjiyad, N., et al., Sugar-free approaches to cancer cell killing. Oncogene, 2011. 30(3): p. 253–64.

27. Greer, S.N., et al., The updated biology of hypoxia-inducible factor. EMBO J, 2012. 31(11): p. 2448–60.

28. MacFarlane, M., G.L. Robinson, and K. Cain, Glucose--a sweet way to die: metabolic switching modulates tumor cell death. Cell Cycle, 2012. 11(21): p. 3919–25.

29. Daskalaki, I., I. Gkikas, and N. Tavernarakis, Hypoxia and Selective Autophagy in Cancer Development and Therapy. Front Cell Dev Biol, 2018. 6: p. 104.

30. Krzywinska, E. and C. Stockmann, Hypoxia, Metabolism and Immune Cell Function. Biomedicines, 2018. 6(2).

31. Majmundar, A.J., W.J. Wong, and M.C. Simon, Hypoxia-inducible factors and the response to hypoxic stress. Mol Cell, 2010. 40(2): p. 294–309.

32. Iurlaro, R. and C. Munoz-Pinedo, Cell death induced by endoplasmic reticulum stress. FEBS J, 2016. 283(14): p. 2640–52.

33. Thomas, L.W., C. Lam, and S.W. Edwards, Mcl-1; the molecular regulation of protein function. FEBS Lett, 2010. 584(14): p. 2981–9.

34. Sathe, P., et al., Innate immunodeficiency following genetic ablation of Mcl1 in natural killer cells. Nat Commun, 2014. 5: p. 4539.

35. O’Sullivan, T.E., et al., BNIP3- and BNIP3L-Mediated Mitophagy Promotes the Generation of Natural Killer Cell Memory. Immunity, 2015. 43(2): p. 331–42.

36. Huntington, N.D., et al., Interleukin 15-mediated survival of natural killer cells is determined by interactions among Bim, Noxa and Mcl-1. Nat Immunol, 2007. 8(8): p. 856–63.

37. Min-Oo, G., et al., Proapoptotic Bim regulates antigen-specific NK cell contraction and the generation of the memory NK cell pool after cytomegalovirus infection. J Exp Med, 2014. 211(7): p. 1289–96.

38. Goh, W., et al., Hhex Directly Represses BIM-Dependent Apoptosis to Promote NK Cell Development and Maintenance. Cell Rep, 2020. 33(3): p. 108285.

39. Donnelly, R.P. and D.K. Finlay, Glucose, glycolysis and lymphocyte responses. Mol Immunol, 2015. 68(2 Pt C): p. 513–9.

40. Keppel, M.P., et al., Activation-specific metabolic requirements for NK Cell IFN-gamma production. J Immunol, 2015. 194(4): p. 1954–62.

41. Mah, A.Y., et al., Glycolytic requirement for NK cell cytotoxicity and cytomegalovirus control. JCI Insight, 2017. 2(23).

42. Chang, C.H., et al., Posttranscriptional control of T cell effector function by aerobic glycolysis. Cell, 2013. 153(6): p. 1239–51.

43. Shin, S., et al., ERK2 Mediates Metabolic Stress Response to Regulate Cell Fate. Mol Cell, 2015. 59(3): p. 382–98.

44. Yao, C.H., et al., Inaccurate quantitation of palmitate in metabolomics and isotope tracer studies due to plastics. Metabolomics, 2016. 12.

45. Yao, C.H., et al., Identifying off-target effects of etomoxir reveals that carnitine palmitoyltransferase I is essential for cancer cell proliferation independent of beta-oxidation. PLoS Biol, 2018. 16(3): p. e2003782.

46. Yao, C.H., et al., Mitochondrial fusion supports increased oxidative phosphorylation during cell proliferation. Elife, 2019. 8.

47. Cheng, T.P., et al., Stability of murine cytomegalovirus genome after in vitro and in vivo passage. J Virol, 2010. 84(5): p. 2623–8.

48. Parikh, B.A., et al., Dual Requirement of Cytokine and Activation Receptor Triggering for Cytotoxic Control of Murine Cytomegalovirus by NK Cells. PLoS Pathog, 2015. 11(12): p. e1005323.

